# Can the brain strategically go on automatic pilot? An fMRI study investigating the effect of if-then planning on behavioral flexibility

**DOI:** 10.1101/2022.11.13.516302

**Authors:** Tim van Timmeren, Nadza Dzinalija, John P. O’Doherty, Sanne de Wit

## Abstract

People often have good intentions but fail to adhere to them. Implementation intentions, a form of strategic planning, can help people to close this intention-behavior gap. Their effectiveness has been proposed to depend on the mental formation of a stimulus-response association between a trigger and target behavior, thereby creating an ‘instant habit’. If implementation intentions do indeed lead to reliance on habitual control, then this may come at the cost of reduced behavioral flexibility. Furthermore, we would expect a shift from recruitment of corticostriatal brain regions implicated in goal-directed control towards habit regions. To test these ideas, we performed a functional MRI study in which participants received instrumental training supported by either implementation or goal intentions, followed by an outcome-revaluation to test reliance on habitual versus goal-directed control. We found that implementation intentions led to increased efficiency during training, as reflected in higher accuracy, faster reaction times, and decreased engagement of the anterior caudate. However, implementation intentions did not reduce behavioral flexibility when goals changed during the test phase, nor did it affect the underlying corticostriatal pathways. Additionally, this study showed that ‘slips of action’ towards devalued outcomes are associated with reduced activity in brain regions implicated in goal-directed control. In conclusion, our behavioral and neuroimaging findings suggest that strategic if-then planning does not lead to a shift from goal-directed towards habitual control.

## INTRODUCTION

At the start of the new year, many people reflect on their future plans and form resolutions. However, they often fail to put their good intentions into practice (Sheeran & Webb, 2016). Strategic ‘if-then’ plans, also known as implementation intentions, are an effective way to support the translation of intentions to actions. For example, instead of formulating an abstract plan such as “I want to lose weight”, an implementation intention links the intended action to a specific cue or situation, e.g. “If I get home, I will eat an apple”, thereby enhancing the probability of success. Indeed, many studies have shown that implementation intentions support behavior change better than goal intentions that merely specify the intended action or outcome (Gollwitzer & Sheeran, 2006). In addition to increasing attention to the relevant cue, the effectiveness of if-then planning has been proposed to rely on creating a strong associative link between the stimulus (S) in the if-part (‘home’) and the response (R) in the then-part (eat an apple), in a manner akin to habits acquired through behavioral repetition (Dickinson, 1985; Thorndike, 1991). These mentally formed S-R associations may allow for automatic action initiation (Gollwitzer, 2014) – a process often referred to as strategic automaticity or ‘instant habits’ (Gollwitzer, 1993, 1999, 2014).

The notion that merely using a verbal action-plan could be sufficient to form a habit is fascinating, because a central assumption in theories of habit formation is that this process critically depends on behavioral repetition. Support for the idea that implementation intentions accelerate habit formation comes from research showing that they increase (self-reported) automaticity (Brandstatter et al., 2001; Orbell & Verplanken, 2010; Parks-Stamm et al., 2007). Therefore, implementation intentions lead to benefits in terms of efficient goal attainment (Gollwitzer, 2014; Gollwitzer & Sheeran, 2006). However, habits developed through behavioral repetition also come at a cost, namely decreased behavioral flexibility (Dickinson, 1985). The question arises, therefore, if the use of implementation intentions also leads decreased flexibility when goals change? This can be investigated using the outcome-devaluation test, an experimental paradigm originally used in rats (Adams & Dickinson, 1981) and later translated to humans (de Wit et al., 2007, 2009; Valentin et al., 2007). In this task, participants first learn to make a response in order to obtain a reward. Subsequently, the value of the outcome associated with that response is devalued, and the ability to flexibly adapt responding to this change in outcome value is measured during an extinction test. Sensitivity to outcome devaluation suggests that behavior is based on knowledge and evaluation of their consequences, and therefore under goal-directed control. If implementation intentions lead to ‘instant habits’ then we would predict reduced sensitivity to outcome devaluation, reflecting a shift from goal-directed towards more rigid, habitual control (Balleine & O’Doherty, 2010; de Wit et al., 2018).

We have previously tested this hypothesis (van Timmeren & de Wit, 2022), using a computerized symmetrical outcome-revaluation task [SORT](Watson, Gladwin, et al., 2022). Participants learn to make a response (Go) to certain ice cream vans to collect valuable ice creams (and points) or to withhold a response (NoGo) to other ice cream vans delivering non-valuable ice creams (and a reduction of (points). To investigate the effect of if-then planning, we instructed them to use verbal implementation intentions for half of the stimuli and use goal intentions for the other half. In the subsequent test phase, some outcome values changed (i.e., outcome revaluation). Whereas participants should continue to respond according to the learnt S-R mappings on value-congruent trials (i.e., still-valuable and still-not-valuable), they should flexibly adjust their behavior on value-incongruent trials (i.e, devalued and upvalued). The results of this previous study suggest that the use of implementation (compared to goal) intentions facilitates instrumental learning, but also impairs performance when some of the signalled outcome values change during the test phase (van Timmeren & de Wit, 2022). This detrimental effect of if-then planning was observed across value-congruent and incongruent trials, suggesting that it was not mediated by strengthened S-R associations (as this would have impacted the value-incongruent trials specifically). Instead, this result may have been driven by reduced goal-directed control. Investigating the neural processes underlying implementation intentions may offer us a window on the underlying (goal-directed versus habitual) processes.

To this end, in the present study we used functional MRI to investigate the neural correlates of if-then planning of instrumental responses on the symmetrical outcome-revaluation task. We capitalized on current insights regarding the neural basis of goal-directed and habitual control to investigate the notion that if-then planning gives rise to ‘instant habits’. Decades of animal research have provided detailed insights into the neurobiology of goal-directed and habitual actions, demonstrating that they are causally supported by anatomically distinct but interacting corticostriatal systems (Balleine, 2019; Balleine & O’Doherty, 2010; Yin et al., 2004). These findings are mirrored by (correlational) neuroimaging evidence in humans, albeit less consistently. Specifically, previous fMRI studies have found that goal-directed control is supported by the ventromedial prefrontal cortex (vmPFC) and caudate while outcome-insensitive habitual actions depend on the premotor cortex and posterior putamen/dorsal striatum (de Wit et al., 2012; Delorme et al., 2016; Morris et al., 2015; Tricomi et al., 2009; Valentin et al., 2007; Watson et al., 2018).

The present study is the first fMRI investigation with the SORT, and we will therefore start with specifying our predictions regarding the general pattern of neural activity independent of intentions. Firstly, we expected that over the course of training (i.e., habit acquisition) activity would increase in regions associated with habitual control while the involvement of regions implicated in goal-directed control would decrease (Liljeholm et al., 2015; Tricomi et al., 2009; Zwosta et al., 2018). Secondly, we expected neural activity during training in these regions to be predictive of revaluation insensitivity in the test phase (de Wit et al., 2009; Liljeholm et al., 2015; Watson et al., 2018; Zwosta et al., 2018). Thirdly, in line with previous work (Valentin et al., 2007; Watson et al., 2018), we hypothesized that in the test phase we would find higher activity in areas implicated in goal-directed action, cognitive control and response conflict when participants flexibly updated their responses and equal (if anything reduced) activity in habit-related regions. Finally, we expected that ‘slips of action’ would be associated with higher activity in habit regions and reduced activity in goal-directed regions (Watson et al., 2018).

Our central aim was to investigate the neural basis of implementation intentions and their effect on behavioral flexibility. To this end, we measured neural activity related to the effect of implementation intentions on acquisition and flexible adjustment of instrumental actions on the SORT. We hypothesized that the use of implementation intentions (compared to goal intentions) during training would lead to increased habit acquisition as reflected by higher accuracy, increased automaticity (measured with the Self-Reported Behavioral Automaticity Index, Gardner et al., 2012) and increased brain activity in habit regions and equal – or if anything reduced – activity in goal-directed regions. Moreover, we expected if-then planning to lead to increased reliance on previously formed S-R associations in the subsequent test phase as indicated by inflexible, habitual responding on value-incongruent compared to value-congruent trials, and higher activity of habit regions during the test phase. Finally, we expected that overcoming mentally rehearsed S-R associations (as part of an if-then plan) would require more goal-directed control and correspondingly engage related neural regions.

## METHODS

All operationalizations, exclusion criteria and main hypotheses and analyses were preregistered on OSF (https://osf.io/yrpxa).

### Participants

Participants were recruited through the participant website of the University of Amsterdam website, flyers and word of mouth. We used the following inclusion criteria: being aged 16-35 years, not having previously participated in a previous study using this same task and any contraindications for MRI. Data collection took place between July and November 2020. Note that this is during the first year of the COVID-19 outbreak; however, no strict lockdowns were implemented during this period in the Netherlands. The study was approved by the Psychology Ethics Committee of the University of Amsterdam and performed in accordance with those guidelines. All participants gave informed consent and received either course credit or financial compensation (15 euro/hour) for their time (total ∼2 hours). An additional €20 voucher was given to the participant with the highest score to motivate participants to perform well on the task.

A total of 47 participants were enrolled, conform our preregistered sampling plan. Our sample size was based on a previous pilot study, which found a significant effect of implementation intentions in 35 participants using the same task and manipulation. Moreover, a power analysis with G*Power (version 3.1.9.3,) showed that our target sample size of n=40 should be sufficient to detect a small behavioral effect (f=0.12) with an α level of .05 and power of .8. Six participants were excluded from all analyses. One participant quit half-way through participation and five participants were excluded based on performance exclusion criteria (see results for details). The remaining 41 participants (22 female, 19 male) had a mean age of 23.2 (SD=4.1) years. All participants had normal or corrected-to-normal vision, and all were right-handed except one who was ambidextrous. All participants were free of neurological or psychiatric disorders and completed or were enrolled in higher professional education at the time of participation, the vast majority being university students. Two participants were native Germans who spoke Dutch fluently, all others were native Dutch speakers.

### Stimuli and materials Procedure

Participants performed a computerized instrumental learning task called the symmetrical outcome revaluation task (Figure 1; Watson, Gladwin, et al., 2022), programmed in Presentation® (version 18.1). Participants played a hungry skateboarder with the objective to collect ice creams to earn points and satisfy their hunger by pressing a response button. The symmetrical nature stems from the inclusion of both valuable and non-valuable outcomes, which and allows comparisons in the test-phase (when outcome values change) between the value-congruent and value-incongruent conditions to be made with the same response-type. See Watson, Gladwin, et al., 2022 for more a more elaborate discussion on the advantages of this task. They were informed that the best performing participant at the end of the study would receive a €20 voucher. Four pictures of ice creams were used: a Cornetto, a Magnum, a Rocket ice lolly and a soft serve ice cream. The task consisted of three phases. First, participants conducted an instrumental training phase without strategic planning outside the scanner, after which they were moved to the MRI scanner and performed an instrumental training phase with strategic planning followed by a test phase (see Figure 1). The total experiment took ∼2 hours, of which 1 hour was spent in the scanner.

**Figure 1.**
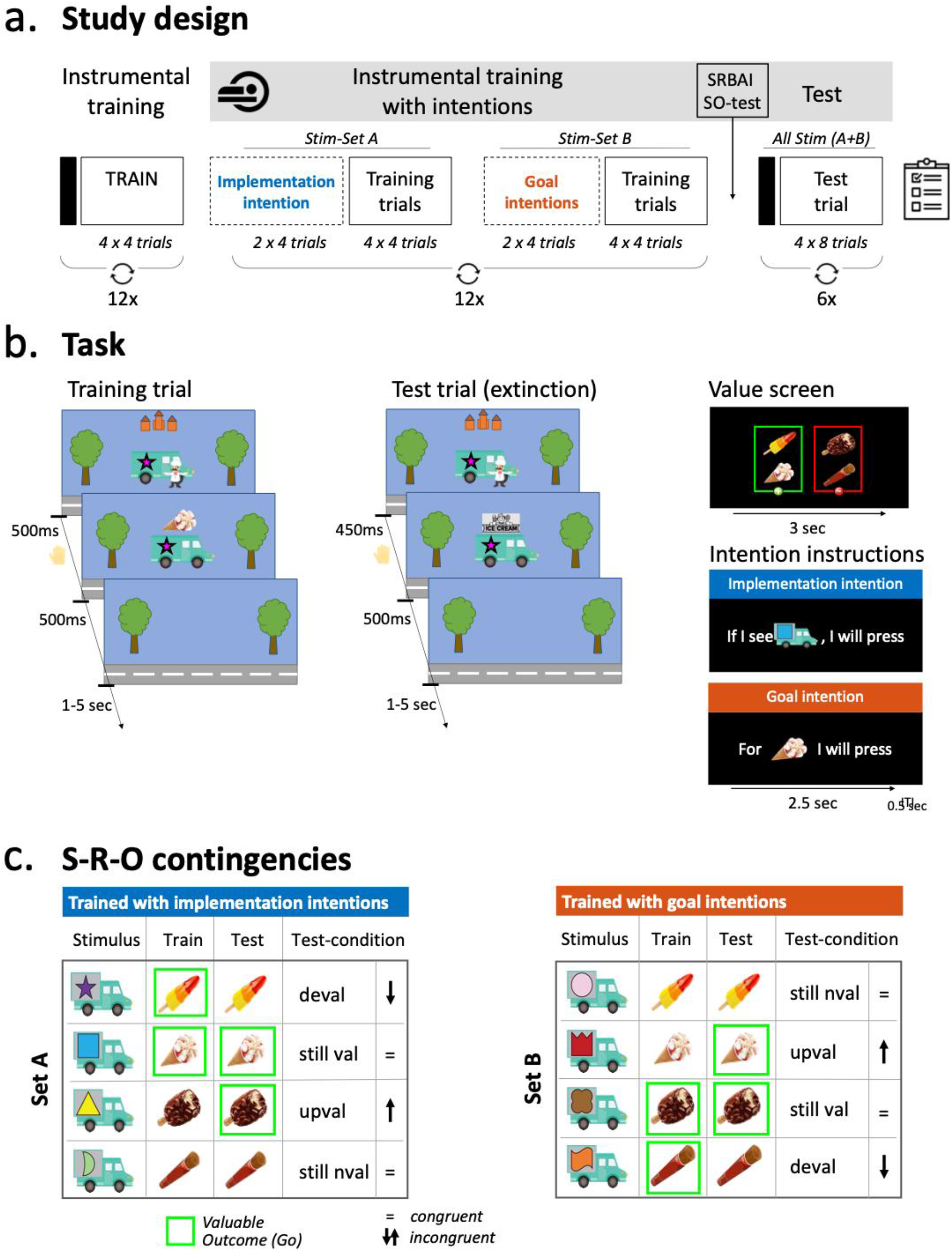
Overview of the study and experimental design. *Note*. Study and experimental design. Participants were told they were playing a hungry skateboarder and their goal was to collect some ice creams and not others to earn points. **a**. Participants first received instrumental training. Each block started with a value-screen (represented by the black rectangle), followed by a block of 16 training trials (see b). Each block contained four vans (pseudo-randomly selected). Training then continued with participants additionally using implementation intentions (trained with van-set A) or goal intentions (trained with van-set B; see c), with intention instructions (see b) being presented before each instrumental learning block. Finally, participants completed six test blocks in which all eight vans (van-set A and B) would appear intermixed and consequently the associated outcome-values of some vans changed compared to training (see c, comparing the ‘Train’ vs ‘Test’ columns). **b**. Train trial: when a van was presented, participants had to decide whether or not to make a response within 500 ms, after which the ice cream appeared (irrespective of a response) on top of the van for 500 ms. Test trial: identical to train blocks but now a banner appeared on top of the van instead of the ice cream, preventing feedback about the outcome (i.e., nominal extinction) and response time was reduced to 450ms. Value screen: the outcome-value screen indicates which ice creams should (in green) and should not (in red) be collected. Intention instructions: vans were trained with either implementation intentions, indicating for which ice cream *van* they should or should not make a response, or goal intentions, indicating for which *ice cream* they should (not) make a response. **c**. An overview of stimulus-outcome contingencies (example set) and associated values across different phases of the task. The contingencies between each ice cream and van remained consistent throughout the whole task, but the *value* of each ice cream (and hence the associated response) was stable only during training. During the critical test phase, the associated outcome values changed (were incongruent) relative to the training value for half of the stimuli (indicated by arrows). This results in four conditions: still-valuable trials (valuable, congruent), upvalued trials (valuable, incongruent), still-not-valuable trials (non-valuable, congruent) and devalued trials (non-valuable, incongruent). For example, the first van always delivered a Rocket, which was valuable throughout training but no longer valuable during test (i.e., devalued). Shown here is an example of the contingencies in one of six test blocks; across the test phase the correct response for each stimulus was equally often congruent and incongruent. Deval = Devalued, still val = still-valuable, upval = upvalued, still not = still-not-valuable trials.

The task used here is almost identical to a previous study in which we tested the same hypothesis behaviorally (van Timmeren & de Wit, 2022), apart from the following changes. To minimize head movements, we used a static version of the task here instead of having ice cream trucks moving across the screen. We added one block of practice with strategic planning before being moved to the scanner, in order for participants to once read the intentions out loud and be able to ask questions. Moreover, we adapted the task to promote S-O learning across intention conditions, to rule out that any effect of implementation intentions on behavioral flexibility would be mediated by reduced contingency knowledge, as was the case in the original behavioral study (REF). To this end, we changed the way in which the blocks were composed in the first part of training (i.e., without intentions): instead of alternating between two sets of four ice cream vans, each block now contains four (out of eight) pseudo-randomly selected stimuli (see *Instrumental training* for details). More than with the block-sets, participants are now forced to pay attention to all outcomes in the value-screen and evaluate for which stimulus they should (not) make a response.

#### Instrumental training

At the start of the task, participants were instructed that their goal was to collect valuable ice creams (which earn points and alleviate hunger) and avoid collecting non-valuable ice creams (which lose points and cause stomach pain) by (not) responding to ice cream vans. There were four different ice creams and prior to each block of instrumental training, participants were shown which two ice creams were valuable (in green) and which two ice creams were not valuable (in red; Fig 1A). The position of the valuable and non-valuable ice creams (left/right) was counterbalanced across participants. Each ice cream was associated with two out of eight vans (Fig 1B): one van always predicting this ice cream as being valuable and the other as being non-valuable. Each block contained only half of the vans: two associated with a valuable ice cream and two with a non-valuable ice cream. Participants were told to find out by trial and error which ice cream truck delivered which ice cream, and that the S-O contingencies would remain the same throughout the whole task. Participants first practiced with different discriminative stimuli (scooters) and outcomes (pizza’s) for two blocks to familiarize them with this procedure. As mentioned previously, the composition of the blocks (i.e., which four out of eight vans were presented during this block) was now pseudo-randomized. The conditions described above allow for six unique combinations of four vans, which were presented twice each (order randomized) during this first part of training for a total of 12 blocks. The contingencies between ice creams and vans and which of the ice creams was valuable/non-valuable were randomized across participants.

Each stimulus was shown four times per block, comprising a total of 16 trials. Trial order was randomized per eight trials, with each van being presented twice in the first and twice in the second half of a block. Each trial started with a jittered 1–5 seconds intertrial interval (ITI). Participants were instructed that they should respond as quickly as possible and before the deliverer disappeared (after 500ms). Irrespective of the response, the associated outcome was then presented for 500 ms. Thus, participants did not receive direct feedback about the accuracy of their response in order to balance the feedback provided for valuable and non-valuable outcomes and to promote goal-directed (R-O) learning and S-O knowledge. Each block ended with a 3-sec feedback screen that displayed accuracy and late responses in that block and total number of points collected (Fig 1D).

#### Instrumental training with intentions

The next phase of training took part in the MRI scanner. Participants were told that instead of seeing which ice creams were valuable or non-valuable, each block would now start with sentences that would help them perform well. These sentences came in two different forms (Fig 1D). *Goal intentions* indicated for each *ice cream* whether or not they should make a response (R-O), formulized as “If I see [picture of an ice cream], then I WILL press”. *Implementation intentions* indicated for each *ice cream van* if they should make a response or not (S-R), formulized as “If I see [picture of an ice cream van] then I WILL (NOT) press”. Each intention was presented for 2500 ms and twice per intention block (randomized order). Half of the stimuli were trained using goal and the other using implementation intentions. Each block of verbal intentions was directly followed by a block of instrumental training (identical to the previous phase) with the corresponding stimuli. Blocks now alternated between two sets of vans, one van-set being trained with implementation intentions (S1-S4, ‘Van-set A’) and one with goal intentions (S5-S8, ‘Van-set B’). Whether the training started with an implementation or goal intention block was counterbalanced across participants. At the end of regular instrumental training and before being moved to the scanner, participants practiced each verbal intention without instrumental training for one block, followed by two blocks (one for each intention type) with instrumental training. During these first few practice blocks outside the scanner, participants were asked to read the intentions out loud. During the subsequent 24 blocks of training with intentions in the scanner, participants were instructed to subvocalize the intentions instead of reading them out loud to minimize head motion. Participants entered the scanner in a head-first supine position and were able to view the screen using a mirror attached to the head coil on which the task stimuli were presented. A button box allowed them to collect ice creams by responding using their right index finger. At the end of training with intentions, participants completed a questionnaire on subjective automaticity (SRBAI) and were tested on their S-O knowledge (details below) (Fig 1E). We had planned to additionally obtain a (pre-intention) baseline measure of these questionnaires, but due to a programming error they were presented *after* the practice blocks with intentions, making them unusable as a baseline measure.

#### Test phase

The test phase was similar to the first training phase (without intentions), but with some important differences. First, as intention blocks were no longer presented, value-screens were again shown at the start of each block, for the duration of 4 sec. Second, participants were informed that the final phase would be more challenging because all ice cream vans would appear intermixed during each block. Therefore, some ice cream vans for which they learned to always make a response during training, now delivered an ice cream that should not be collected and so they should withhold making a response (and vice versa). We instructed them to pay extra attention to the value-screens because some ice creams vans that previously always delivered ice creams would now lead to a reduction of points (i.e., devalued trials) and vice versa (i.e. upvalued trials). Third, participants were told that the ice cream deliverers placed a banner on top of their van, blocking the view of the ice cream they delivered (i.e., nominal extinction). But because each van still kept on delivering the same ice cream as during training, they should base their choice on what they learned before. Finally, the feedback screens presented at the end of each block no longer included information on the *accuracy* of their responses, but only the percentage of responses, non-responses, and late responses. We did this to prevent outcome-based learning during the test phase. We explicitly instructed participants that each block contained an equal amount of valuable and non-valuable outcomes so they knew they should aim for a 50%/50% distribution. Participants completed a total of six test blocks.

An important consequence of each block containing all eight stimuli is that some vans would now deliver an ice cream with a value *incongruent* with the value during training. This resulted in four different conditions for each intention-type: ‘still-valuable’ – the outcome signaled by this stimulus was always valuable during training and also valuable during this test block (i.e., congruent); ‘upvalued’ – the outcome signaled by this stimulus was not valuable during training but valuable during this test block (i.e. incongruent); ‘still-not-valuable’, – the outcome signaled by this stimulus was not valuable during training and still-not-valuable during this test block (i.e., congruent); and ‘devalued trials’ – the outcome signaled by this stimulus was always valuable during training but not valuable during test (i.e., incongruent).

We also shortened the response window to 450ms to force rapid responding, which has been shown to boost the expression of habitual slips (Hardwick et al., 2019). However, because a lot of participants responded just after the 450-ms time limit, we decided to include responses up to 600ms for both the behavioral and fMRI analysis to increase the number of included trials in the fMRI analyses. This change did not significantly impact the pattern of behavioral results, which was unsurprising as the test phase was conducted in extinction, meaning that no performance feedback was provided during this period.

### Self-Report Behavioral Automaticity Index (SRBAI)

The SRBAI (Gardner et al., 2012) is a four-item scale that captures self-reported habitual behavior patterns that we adapted for to assess automaticity for (not) responding to the ice cream vans. Participants were presented with each ice cream van and asked to indicate the associated response (press or not press) and the degree to which (not) making a response was something they did: “automatically”, “without having to consciously remember”, “without thinking” and “before I realize I am doing it”. Each item was scored on scale ranging from 1 (strongly disagree) to 100 (strongly agree). The SRBAI scale was previously shown to have good reliability and validity (Gardner et al., 2012). Before the four SRBAI items appeared, participants were asked to indicate which response was associated with that stimulus (“making a response” / “not making a response”) to test S-R knowledge. Cronbach’s alpha was calculated separately for each of the four conditions (2 intentions x 2 values), using the 8 test items (four SRBAI questions for the two stimuli per condition). The results indicate high internal reliability, with alpha ranging from .91-.95. The final score was calculated separately for each intention by taking the mean across the 4 items (range: 1-100), with higher scores reflecting more automatic behavior.

### Test of stimulus-outcome knowledge

Participants were asked about their knowledge of the S-O contingencies by asking them for each ice cream vans which ice cream it delivered. After selecting one of the four ice creams, participants were asked to indicate how confident they were about their decision (0-100). Composite scores, reflecting S-O knowledge, were calculated for each intention and separately for Go- and NoGo-trained stimuli by multiplying percentage of correct S-O contingencies (0/50/100%) with percentage mean confidence.

### Pre-registered behavioral data analysis

Behavioral data analyses were performed using IBM SPSS Statistics 25 for Mac (IBM, New York, NY, USA) for frequentist statistics and JASP version 0.16.3 (JASP Team, 2018) for Bayesian statistics. For data analysis purposes the training data were collapsed across blocks of three, referred to as block-sets. Accuracy is reflected by the percentage of trials on which a correct response was made, calculated by the number of correct responses divided by the total number of trials. In line with the fMRI analyses, trials on which a late response was made were not included in this analysis. To assess that learning took place over the first part of the training without intentions, accuracy was analyzed using a 2×4 repeated measures analysis of variance (ANOVA) with within-subject factors value (valuable or non-valuable) and block-set (1-4). The second part of training was analyzed using a 2×2×4 repeated measures ANOVA, with intention type (implementation or goal-intention) as an additional factor. Reaction times for correct responses were analyzed with similar ANOVAs, but for valuable trials only.

For the test phase, data were analyzed using a 2×2×2 repeated measures ANOVA with three factors: intention-type (implementation or goal-intention), test value (valuable or non-valuable during test) and congruency (congruent or incongruent with value during training). Thus, for each intention-type there are four conditions: still-valuable trials (valuable, congruent), upvalued trials (valuable, incongruent), still-not-valuable trials (non-valuable, congruent) and devalued trials (non-valuable, incongruent). Again, reaction times were analyzed using similar ANOVAs. Note that eight participants were excluded from the NoGo analyses because they performed perfectly on still-not-valuable trials and thus did not make any response.

Subjective automaticity (SRBAI scores) for responding to stimuli trained with implementation and goal intentions at the end of training was compared using a paired *t*-test. Finally, the relationship between automaticity and the ‘revaluation insensitivity’ index was tested for both intention types separately using correlational analyses. A revaluation insensitivity index was calculated for each intention type by taking the difference between accuracy for congruent and incongruent test trials separately for Go (still-valuable minus devalued) and NoGo-trained stimuli (still-not-valuable minus upvalued), with higher revaluation insensitivity scores indicating more habitual performance. Kendall’s tau was used as the four revaluation indices and SRBAI scores were not normally distributed.

In the case of violations of sphericity, we report Greenhouse–Geisser corrected degrees of freedom and p-values. In addition to 95% confidence intervals, partial eta squared (*η*_p_^2^) for the ANOVAs and Cohen’s d for paired t-tests are reported as estimates of effect sizes. As preregistered, we additionally conducted corresponding Bayesian analyses for the main analyses of interest and report corresponding Bayes Factors. For key null results (p > 0.05) we report Bayes Factor_01_, which quantifies the relative evidence in favour of the null hypothesis H0 over the alternative hypothesis (H1). For key significant results (p < 0.05) we report the Bayes Factor_10_, which quantifies evidence in favour of the alternative hypothesis over H0 (and is identical to 1/BF_01_). Bayes factors were interpreted according to (Wetzels et al., 2011, Table 1), with Bayes Factors between one and three reflecting anecdotal support, Bayes Factors larger than three reflecting substantial support and Bayes Factors larger than ten reflecting strong support. In all Bayesian analyses, JASP’s default priors were used.

**Table 1.** Imaging results of the training phase (exploratory analyses)

| Contrast | Region | MNI coordinates<br>(x,y,z) |  |  | Cluster size<br>(voxels) | z-score<br>at peak<br>level | Correction |
| --- | --- | --- | --- | --- | --- | --- | --- |
| Increase over training<br>blocks (Go) | Caudate Nucleus head | 22 | 6 | 30 | 443 | 4.37 | Cluster |
|  | Amygdalo-hippocampal<br>junction | -10 | -4 | -14 | 348 | 5.17 | Peak |
|  | Angular gyrus | 20 | -52 | 38 | 214 | 4.92 | Peak |
|  | Posterior Putamen | 26 | -20 | 4 | 34 | 3.96 | SVC Tricomi |
| Decrease over training<br>blocks (Go) | Anterior caudate L | -24 | 10 | 2 | 912 | 6.53 | Cluster |
|  | Anterior caudate R | 24 | 10 | -4 | 537 | 6.39 | Cluster |
|  | Primary motor/SMA | 8 | -24 | 60 | 860 | 5.44 | Cluster |
|  | Hippocampus/putamen | 43 | 14 | -8 | 657 | 4.83 | Cluster |
|  | Temporal cortex L | -46 | -46 | -4 | 591 | 5.69 | Cluster |
| Goal > implementation<br>intentions block-set 1<br>(Go) | Anterior caudate | 13 | 18 | -4 | 40 | 3.69 | SVC striatum |

*SMA=Supplementary Motor Area; SVC=Small Volume Correction; L=left, R=Right*

### MRI data acquisition

All magnetic resonance imaging (MRI) was performed on a 3 Tesla, full-body Achieva dStream MRI-scanner (Philips Medical Systems, Best, The Netherlands) equipped with a 32-channel head coil. After entering the scanner, a low-resolution survey scan was made to determine the location of the field-of-view.

Functional MRI scans were acquired at a ∼30° angle from the anterior-posterior commissure line to maximize signal sensitivity in orbital regions (Deichmann et al., 2003) using a T2*-weighted single-shot gradient echo imaging sequence with the following parameters: repetition time (TR) = 2000 ms; echo time (TE) = 28 ms; flip angle = 76.1°; voxel size = 3 mm^3^ with 0.3 mm slice gap; matrix size = 80×78; number of slices = 36; field of view (FOV) = 240 × 118.5 × 240 mm. The training with intentions was split in two runs of 598 scans each, while a total of 415 scans were acquired for the test phase. The first six volumes of each run were discarded to allow T1 saturation to reach equilibrium.

A high-resolution T1-weighted structural image was acquired before the final run (while participants completed the post-training SRBAI and SO-test) using an MPRAGE sequence with the following parameters: voxel size = 1 mm^3^; FOV = 240 × 220 × 188 mm; TR = 8.2 ms; TE = 3.7 ms, 220 slices, flip angle = 8°.

### fMRI data analysis

#### Image preprocessing

MRI data were first converted to Brain Imaging Data Structure (BIDS) format using in-house scripts. An initial check of data quality was done by visually inspecting the image-quality metrics derived from MRIQC v0.15.0 (Esteban et al., 2017). Data was preprocessed using fMRIPrep v20.1.1 (Esteban et al., 2019) with the default processing steps. These essentially included brain extraction, segmentation, and surface reconstruction of the structural T1 image; spatial normalization of both the structural and functional data to MNI space; and head motion estimation, coregistration, susceptibility distortion correction and resampling to 2 mm^3^ of the functional data. No slice-timing correction was performed. A comprehensive description of the preprocessing pipeline is available from the citation boilerplate (See Supplementary analysis).

#### fMRI statistical analyses

The preprocessed functional data was further analyzed using Statistical Parametric Mapping software (SPM12, Wellcome Trust Centre for Neuroimaging, London). The data was spatially smoothed using a Gaussian kernel with a full width at half maximum of 8mm and all functional data was high pass filtered (with a 128 sec cutoff) to remove slow signal drifts.

##### First-level analysis

For the first-level analysis of the fMRI data, a general linear model (GLM) was constructed for each subject, concatenated over all three runs from the training and test phase. For the data on training with intentions, trial onsets of valuable stimuli and non-valuable stimuli for implementation and goal intentions were modeled using stick functions, making a total of 4 conditions. To look at the effect of time on training, these were modeled as separate regressors per 3 blocks, making a total of 4 training block-sets. Only correct trials (i.e., where an accurate (non)response was made) were included. Blocks of verbal rehearsal of implementation and goal intentions were additionally modeled as blocks of 28 sec (total duration of eight 3.5 sec trials). For the test phase, stick functions modeled the trial onsets of still-valuable and still-not-valuable (‘value-congruent’; the outcome value is congruent with training phase) and devalued and upvalued (‘value-incongruent’; the outcome value is not congruent with training phase) stimuli that were trained with implementation or goal intentions separately, making a total of 8 regressors. To investigate BOLD activity during habitual (c)omission errors (habitual ‘slips’ in case of incongruent trials), separate regressors were included for incorrect trials for all conditions. The following regressors of no interest were included separately for each run: one regressor for errors (only for training, as test-errors/’slips’ were modelled as regressors of interest) and late trials, keypresses, feedback-displays, value-screens (only for test phase), and six realignment parameters capturing rotation and translation to correct for residual subject motion. Three session constants were included in the model. All onsets were then convolved with the canonical hemodynamic response function and an autoregressive AR(1) model was used to correct for serial correlations. The GLM was regressed against the fMRI data to generate parameter estimates for each subject.

Regressor-specific first-level contrast images were created for the training- and test-regressors modeling the different conditions of interest to construct the planned second-level full factorial models. These contrasts of parameter estimates were then entered into between-subjects analyses of variance (ANOVAs) to generate group-level random-effects statistics. To test for a difference in learning between intention types, contrasts of parameter estimates of the instrumental training phase were entered into a 2 × 4 × 2 (value by block-set by intention) factorial ANOVA. Following estimation of the second-level model, t-tests were specified by adding linear weights to each instrumental training block-set, modeling increases over training as [-1.5 -0.5 0.5 1.5] and decreases as [1.5 0.5 -0.5 -1.5].

Additionally, first-level contrast images were created. To assess the effect of planning during training, contrasts were created comparing training with implementation versus goal intentions (across all blocks, separately for Go and NoGo trials). To examine markers of goal-directed control during test, we compared correct *congruent* trials with correct *incongruent* trials (i.e. [still-valuable Go > upvalued Go] and [still-not-valuable NoGo > devalued NoGo]). We also investigated situations where participants fail to adapt to the new outcome value and continue to respond according to the learned S-R association by comparing *incorrect* incongruent trials (i.e., ‘slips of action’) with *correct* incongruent trials. Again, separate contrasts were created for test-Go- and test-NoGo trials (i.e., [devalued Go > upvalued Go] and [upvalued NoGo > devalued NoGo]). Finally, we also created a similar contrast comparing incorrect incongruent trials (slips) with *correct congruent* trials (i.e., [devalued Go > still-valuable Go] and [upvalued NoGo > still-not-valuable NoGo]). More information about the rationale behind these contrasts is provided in the Results section. To assess the effect of planning strategy on test performance, the same test-phase contrasts were constructed but looking for an interaction with intention type (e.g. [still-valuable Go > upvalued Go X implementation > goal intention]). Parameter estimates generated from these first-level analyses were entered into a random-effects group analysis, and linear contrasts were used to identify significant effects at the group level.

In our preregistration, we identified several regions of interest (ROIs) for habitual control, goal-directed control, response conflict and implementation intentions. Three separate masks were created based on these ROIs to apply small volume correction (SVC). Apart from a striatal ROI (encompassing the bilateral caudate, putamen and NAcc from the AAL atlas (Tzourio-Mazoyer et al., 2002)), however, applying SVC with the three preregistered ROIs did not alter the pattern of results. This may be due to the large number of regions included in the ROIs (especially the goal-directed ROI) which limited the effect of small-volume correction, because using such a large mask does not significantly increase power. Therefore, we have opted to report the whole-brain confirmatory analyses.

Whole-brain t-maps (without thresholding) of the main fMRI contrasts are available at https://neurovault.org/collections/13191/.

## RESULTS

All analyses reported in this section were pre-registered at the start of this study, unless indicated otherwise in the text. We generally followed the pre-registered analysis plan, but in some cases the results prompted us to further explore the data. We should also point out that we preregistered these hypotheses before finishing data analysis of our related behavioral study (van Timmeren & de Wit, 2022). Hence, we preregistered the same behavioral hypotheses for this study, even though the original behavioral study only partially supported our initial predictions – a point we will come back to in the discussion. We therefore incidentally deviate from the preregistration to keep our analyses in line with analyses and findings from the behavioral study, which is always clearly indicated.

The total final sample used for the analyses consisted of 41 participants, after excluding the following participants. Based on the preregistered exclusion criteria, no participants were excluded on the training criterion (<80% accuracy in last block-set of training), while three were excluded because they made <25% responses on upvalued trials trained with goal intentions in the test-phase. The goal of this criterion was to ensure that participants understood the test-phase instructions and updated their performance accordingly, while not excluding participants based on the manipulation of interest (i.e., implementation intentions). We additionally excluded 2 participants (post hoc) based on a very low overall response rate during the test phase. Although these participants made (just) >25% upvalued responses, we deviated from the preregistration because they were outliers on the overall response rate and responded on less than 1 out of 3 trials during the test, despite receiving explicit instruction to aim for a response rate of ∼50% and receiving feedback about that at the end of each block. Hence, they did not follow the test-phase instructions and their performance is not reliable. Note that this criterion is independent of actual task performance (accuracy) and that the in-/exclusion of these two participants does not change the general pattern of behavioral nor fMRI results.

### Behavioral results

#### Training phase without intentions

As expected, participants learned to make correct responses over the first part of training, as revealed by a significant main effect of block-set on accuracy (*F*_2.46, 98.20_=16.74, p<0.001 *η*_p_^2^=0.30) and a marginally significant effect of block-set on reaction time (*F*_2.45, 98.07_=2.75, p=0.058). Participants learned to make Go and NoGo responses equally well (*F*_1, 40_=2.00, p=0.17, *η*_p_^2^=0.05).

#### Instrumental training with goal versus implementation intentions

Following the first 12 blocks of instrumental training without planning, intentions were introduced during a practice block outside the scanner. Although we did not preregister to analyze those data, for completeness and in line with our previous behavioral study with this paradigm investigating the same question (van Timmeren & de Wit, 2022), we conducted a paired t-test comparing the final block of training without intentions to the practice block. This analysis revealed that participants benefitted from if-then planning on the valuable Go trials, as reflected by higher accuracy (M=96.1, SD=12.4) relative to the preceding (pre)training block-set (baseline: M=91.8, SD=9.1, z_40_ = -2.57, *p*=0.01, d=-.59, 95% CI = [-.81 – -.22]), while reaction times were not affected (*t*_40_ = .01, *p*=0.99, d=.001). In contrast, the use of goal intentions negatively impacted both accuracy (M=87.6, SD=14.7, z_40_=1.86, p=0.065, d=.34, 95% CI = [.01 – .69]) and reaction times (*t*_40_ = -2.03, *p*=0.049, d=-.32) compared to (pre)training. No significant effects of intention were seen for NoGo trials (all p>.31).

Subsequently, when instrumental training was resumed during the scanning session, the advantage of if-then planning was initially still apparent on valuable Go trials. In addition to a strong main effect of value, driven by participants performing better overall on valuable compared to non-valuable trials (*F*_1,84.47_=10.93, p=.0002, *η*_p_^2^=0.22), we found the expected preregistered 3-way interaction between intention, value and block-set (*F*_3,103.14_=6.45, p<0.001, *η*_p_^2^=0.14). Separate analyses of valuable and non-valuable trials revealed a significant intention–by–block interaction for valuable (*F*_3,81.78_=6.21, p=0.003, *η*_p_^2^=0.13), but not for non-valuable trials (*F*_3,120_=1.88, p=0.14, *η*_p_^2^=0.05). The significant effect on the valuable Go trials was driven by higher accuracy with implementation compared to goal intentions during the first block-set (z_40_ = 3.34, p<0.001, d = .85, 95% CI = [.64 – .94]). At the end of training (block-set 4), there was no longer a significant effect of intention type on accuracy (z_1,40_=-.34, p=.80, *η*_p_^2^=-1.43).

The analysis of reaction times revealed a main effect of intention type (*F*_1,40_=12.08, p=.001, *η*_p_^2^=0.23), with faster responses during blocks trained with implementation intentions (Median=365ms, SD=17) compared to goal intentions (Median=374ms, SD=20), and a marginal effect of block-set (*F*_2.4,98.6_=2.31, p=.09, *η*_p_^2^=0.05), but no significant interaction.

##### Symmetrical outcome-revaluation test

As expected, learned S-R associations had a clear impact on performance during the test phase (Figure 2B), as revealed by a main effect of congruence (*F*_1,40_=65.08, p<.001, *η*_p_^2^=0.62). Because test value showed significant interactions with both congruence (*F*_1,40_=10.73, p=.002, *η*_p_^2^=0.21) and intention type (*F*_1,40_=5.94, p=.02, *η*_p_^2^=0.13), separate follow-up comparisons were conducted for Go (associated with still-valuable and upvalued outcomes) and NoGo (associated with still-not-valuable and devalued outcomes) trials. Main effects of congruence were seen for both the Go (*F*_1,40_=16.82, p<.001, *η*_p_^2^=0.30) and NoGo (*F*_1,40_=56.46, p<.001, *η*_p_^2^=0.59) stimuli. As can be seen in Figure 2B, the congruency effect was larger for NoGo trials mainly due to participants struggling more on devalued trials, where they had to suppress responding to discriminative stimuli that previously signaled a valuable outcome. Importantly, we were interested in the effect of implementation intentions on test performance. Firstly, frequentist analysis of the Go test trials suggested that performance was worse when trained with implementation compared to goal intentions (*F*_1,40_=5.48, p=.02, *η*_p_^2^=0.12), but Bayesian statistics showed that this evidence was inconclusive (Bayes Factor_10_=1.49). Importantly, in contrast to our preregistered hypothesis, there was no evidence for reduced flexibility as a consequence of if-then planning: the expected interaction of congruence with intention type failed to reach significance (*F*_1,40_=1.52, p=.23, *η*_p_^2^=0.04, Bayes Factor_01_=0.85). Given the direct relevance of the comparison between intentions for our research question, we followed these analyses up with separate (exploratory) paired t-tests for still-valuable and upvalued trials in order to also report Bayesian evidence against a difference. Findings indicate that intentions only had a significant negative effect on (congruent) still-valuable (t_40_=-2.66, p=.01, Bayes Factor_10_=3.68) but not on (incongruent) upvalued trials (t_40_=-.75, p=.46, Bayes Factor_01_=4.54). Finally, for the NoGo stimuli (still-not-valuable and devalued), no main (*F*_1,40_=.42, p=.52, Bayes Factor_10_=5.36) nor interaction (*F*_1,40_=.06, p=.81, Bayes Factor_01_=5.57) effects of intention type were observed. We also analyzed reaction times during the test phase. A value x congruence interaction (*F*_1,32_=49.47, p<.001, *η*_p_^2^=0.61) prompted separate analyses for trials trained with Go responses (still-valuable and devalued) and for trials trained with NoGo responses (still-not-valuable and upvalued). Interestingly, we found significantly faster reaction times on devalued trials (M=418ms, SE=8.8) relative to still-valuable (M=443ms, SE=6.8; *F*_1,40_=12.56, p=.001, *η*_p_^2^=0.24), in line with the idea that habitual slips of action are triggered fast and efficiently before one has the chance to suppress them. As late responses were excluded from this analysis, we ran an additional analysis including reaction times for late responses to make sure that this effect was not driven by a higher number of (excluded) late responses on devalued trials. This analysis showed an even stronger main effect of congruence (*F*_1,40_=14.84, p<.001, *η*_p_^2^=0.27). No other significant effects of reaction times were found (all p>.22).

**Figure 2.**
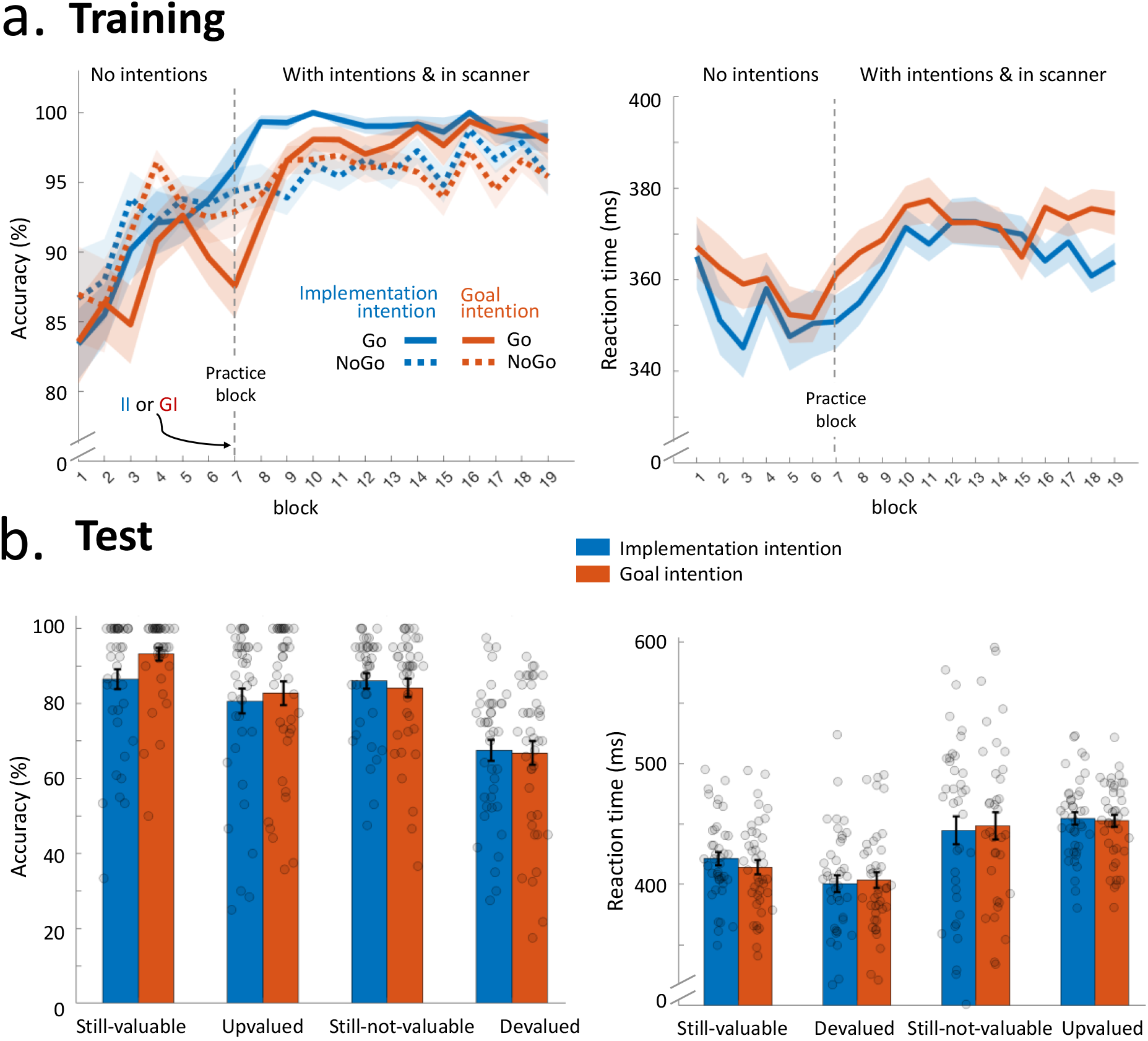
Behavioral results *Note*. **a**. Over the course of training, participants learned to successfully respond for stimuli associated with valuable outcomes (Go) and to withhold making a response for stimuli associated with non-valuable outcomes (NoGo), as reflected by increasing accuracy rates. After six blocks of regular training, some stimuli continued to be trained using implementation intentions (blue) while others were trained with goal intentions (blue). Following one block of practice (black dotted line), participants were moved to the scanner and resumed training with intentions. Accuracy was significantly higher initially when using implementation intentions, but towards the end of training performance was almost perfect for both implementation and goal intentions. Across training with intentions, participants were faster during blocks trained with implementation versus goal intentions. **B**. During the test phase, for some stimuli the associated outcome changed in value (and thus response) compared to training (upvalued and devalued; see Figure 1C) and participants had to flexibly update their responses accordingly. For other stimuli, the associated value and response remained congruent with training (still-valuable and still-bot-valuable). Participants responded less accurately for incongruent compared to congruent trials, reflecting inflexibility as a consequence of learned S-R contingencies during training. However, training with implementation intentions did not lead to reduced flexibility. Similarly, there was no significant effect of training with implementation intentions on reaction time. (Shaded) error bars represent standard error of the mean. II = implementation intentions; GI = goal intentions.

##### Self-reported automaticity and S-O knowledge

Self-reported automaticity was at a high level overall (median 80.4%, SD=16.7), but did not differ between intentions (t_40_=-.98, p=.34), nor did subjective automaticity correlate with revaluation insensitivity for implementation (r_τ_=-.09, p=.57) or goal intentions (r_τ_=.22, p=.17).

Following van Timmeren & de Wit (2022), we also explored differences in S-O knowledge between intention types and their relationship with overall test accuracy. S-O knowledge was high (median=89.8%, SD=22.1) and, contrary to our previous study, no longer differed significantly between intention types (p=.16), values (p=.07), or their interaction (p=.35), suggesting that the adaptation we made to the task had the desired effect (i.e., using a pseudo-random selection of stimuli instead of alternating between two block-sets in the first part of training, see Methods). S-O knowledge did correlate positively with overall test accuracy for both implementation intentions (r_τ_=.30, p=.008, 95% CI = [.08, .52]) and goal intentions (r_τ_=.39, p<.001, 95% CI = [.21, .57]).

## Conclusions behavioral results

We provide evidence for habit learning, as indicated by the general effect of previously learned S-R mappings on the ability to flexibly adapt responding when the cue signals a revalued outcome (i.e., incongruent). Importantly, while if-then planning seemed to increase efficiency relative to goal intentions, as reflected in superior acquisition, this was not at the expense of flexibility when outcome values changed in the test phase.

### Neuroimaging results

#### Instrumental training: across intentions (exploratory)

First, we were first interested to explore general learning effects across intention types because this was the first time the SORT was used in the fMRI scanner. These analyses showed that over the course of Go training (i.e., on valuable trials), activity increased linearly in the head of the caudate nucleus extending into ACC, the left amygdalo-hippocampal junction, and the angular gyrus (all p<0.05 FWE corrected, Table 1). In this same contrast we also observed a cluster in the posterior putamen, which survived a small-volume correction for the posterior putamen ROI (i.e. p_FWE_<0.05 with SVC, defined as a 10-mm sphere at peak value of the cluster that showed a significant increase over training in the study of Tricomi et al. (2009); x = 33, y = -24, z = 0). On the other hand, activity decreased over training in the bilateral anterior caudate (a more ventral part of the striatum), primary motor cortex (extending to mid-posterior cingulate), hippocampus extending into the putamen, and the left temporal cortex (all p<0.05 FWE corrected). In contrast, on NoGo trials there were no voxels that showed a significant linear change over training blocks.

##### Instrumental training: comparing goal and implementation intentions

We then examined whether strategic planning affected instrumental training. The contrast comparing the average BOLD signal of trials trained with implementation intentions and goal intentions did not reveal any significant activations, neither on Go nor NoGo trials. We also tested for differences in learning between intentions over the course of training by adding linear weights to block-sets to compare increased activity over block-sets during implementation intentions with decreased activity during goal intentions, and vice versa. However, both tests of this interaction failed to show significant differences.

The finding that implementation intentions showed the most pronounced effect behaviorally *early* in training prompted us to conduct an *exploratory* analysis of only the first training block-set. This analysis revealed significantly decreased activation in the anterior caudate (p<.05 familywise error (FWE) rate small-volume corrected, z=3.69) on trials trained with implementation intentions compared to goal intentions (Figure 3A and Table 1). For visual purposes, the extracted average BOLD signal from the anterior caudate cluster is shown separately for each block-set and intention in Figure 3B. As can be seen here, activity was indeed lower on implementation intention trials during the first block-set only, and subsequently decreased for both intentions. We also found the same contrast to show decreased activity for implementation relative to goal intentions at an uncorrected threshold in the left insula (*z*=3.71; *x*=-42, *y*=20, *z*=0) and the right lateral orbitofrontal cortex (OFC) (p_FWE_=.061, z=4.25; *x*=26, *y*=59, *z*=14). However, because these results did not survive cluster correction, we refrain from interpreting them further. To rule out that these findings were driven by reaction times, which were significantly shorter for implementation compared to goal intentions, we performed an additional analysis controlling for trial-by-trial reaction time by including a parametric regressor (one for each of the two training runs) with reaction times for each trial. This had no significant impact on the results, and we could qualitatively replicate all reported findings.

**Figure 3.**
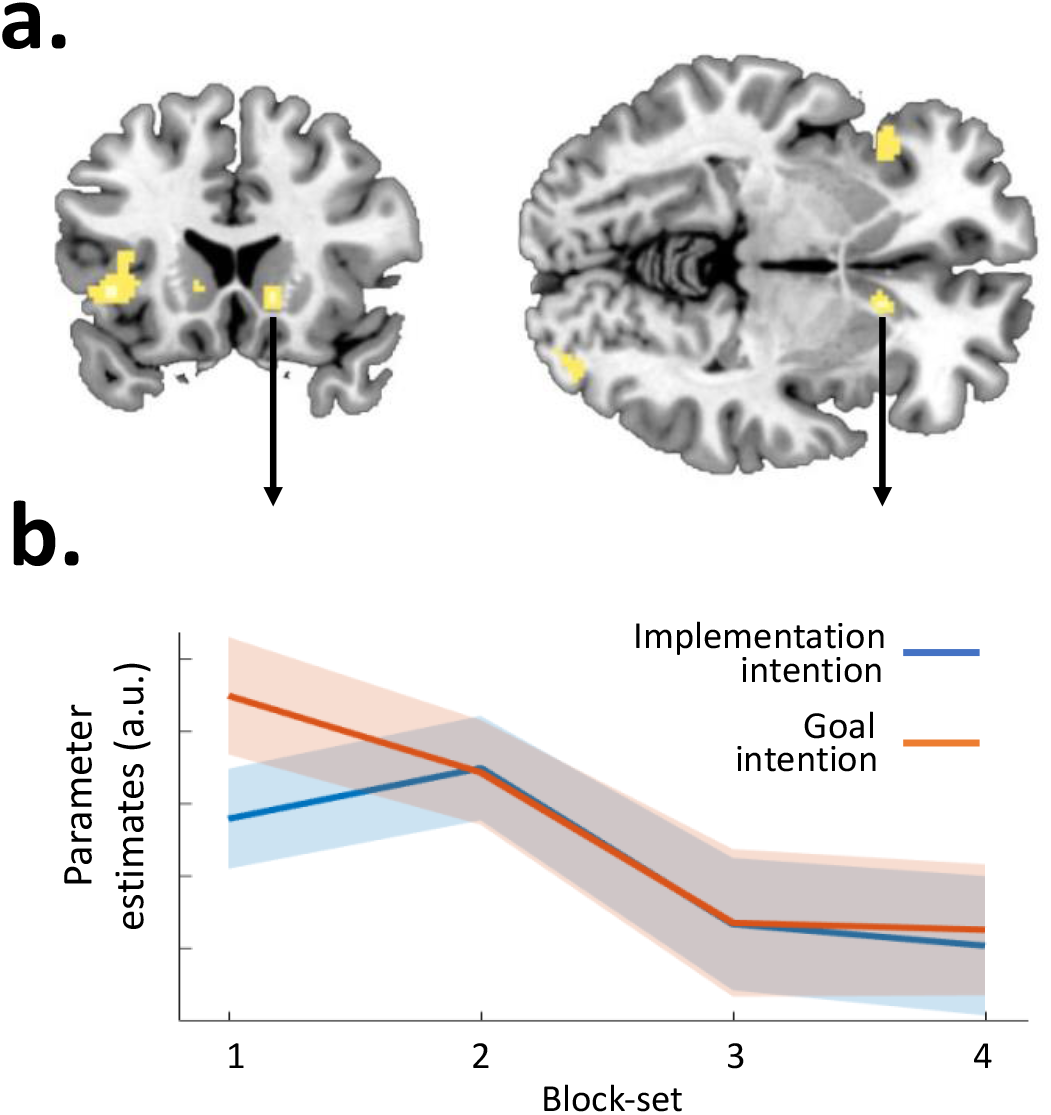
Lower activity in the right anterior caudate early in training for implementation compared to goal intentions. *Note*. **A**. Voxels that showed significantly lower activation during the first block-set of training with implementation compared to goal intentions on Go-trials (shown at p<.001 uncorrected). **B**. parameter estimates extracted from this anterior caudate cluster (peak at x=13, y=18, z=-4) over block-sets. Error-bars represent 95% confidence intervals. a.u.: arbitrary units.

##### Neural predictors of test performance

In order to determine whether brain activity during instrumental training with implementation intentions was predictive of test performance, we tested whether the average BOLD signal during training covaried with the revaluation insensitivity score. This preregistered test did not reveal significant neural predictors of test performance. For completeness we also exploratively ran this analysis separately for goal intentions and across intentions, but this similarly did not reveal any significant results.

##### Symmetrical outcome-revaluation test: markers of goal-directed versus habitual performance

In the test phase, changes in outcome value create conflict between goal-directed control and learnt S-R associations. Specifically, in order to perform the correct response on incongruent trials (i.e., upvalued Go and devalued NoGo), participants have to exert goal-directed control and override the learnt S-R mapping. Conversely, on congruent trials (still-valuable Go and still-not-valuable NoGo), participants can rely on the learnt S-R associations. The advantage of the symmetrical outcome-revaluation test (compared to the original slips of action test) is that we can compare congruent and incongruent trials with each other unconfounded by test outcome value (and therefore required response: i.e., Go or NoGo). Therefore, to examine markers of goal-directed control, we firstly compared upvalued Go with still-valuable Go responses and found that this was associated with increased right insula activity (p<0.05 FWE corrected, z=4.16; Table 1). No significant activations were seen in the contrast between devalued NoGo and still-not-valuable NoGo trials.

To identify regions where participants fail to adapt and continue to respond according to the learned S-R association, we contrasted incorrect incongruent trials (devalued Go and upvalued NoGo) to correct incongruent trials (upvalued Go and devalued NoGo, respectively), as the latter arguably require most goal-directed control to override the learnt S-R mapping. The contrast comparing devalued Go responses (i.e., slips of action) with upvalued Go responses is shown in Figure 4A, and revealed increased activity in a fronto-parietal network, including the left anterior insula extending to the inferior lateral prefrontal cortex, supplementary motor area, dorsal anterior cingulate cortex, bilateral inferior parietal lobule and supramarginal gyrus (all p<0.05 FWE corrected, Table 1). Conversely, lower activity during slips of action compared to upvalued Go responses was seen in the left anterior cingulate cortex extending into caudate nucleus, premotor/primary motor cortex, left lateral OFC, bilateral superior parietal lobe and several occipital/primary visual areas (all p<0.05 FWE corrected, Table 1). While the previous contrast between devalued slips and correct upvalued Go responses maximizes the difference between habitual versus goal-directed control, the conditions differ in terms of the original training outcome value(as well as test value). To mitigate this, we proceeded to compare devalued slips to still-valuable Go responses, which only differ in their test outcome value. Thus, this contrast compares trials on which participants correctly continued responding according to the learned S-R association with trials on which they failed to override this association. Although we have used the exact same approach previously (in the study of Watson et al., 2019, the ‘slips versus respond valuable’ contrast), this contrast was not preregistered, and should thus be considered exploratory. Similar to the comparison with upvalued Go responses, this comparison of slips with still-valuable Go responses revealed increased anterior insula activity (bilaterally) during slips, but decreased activity in vmPFC (extending to NAcc), primary motor cortex, paracentral lobule, a large occipital cluster and large parietal clusters (bilateral) including the angular gyrus and inferior parietal lobule (all p<0.05 FWE corrected, Figure 4A).

**Figure 4.**
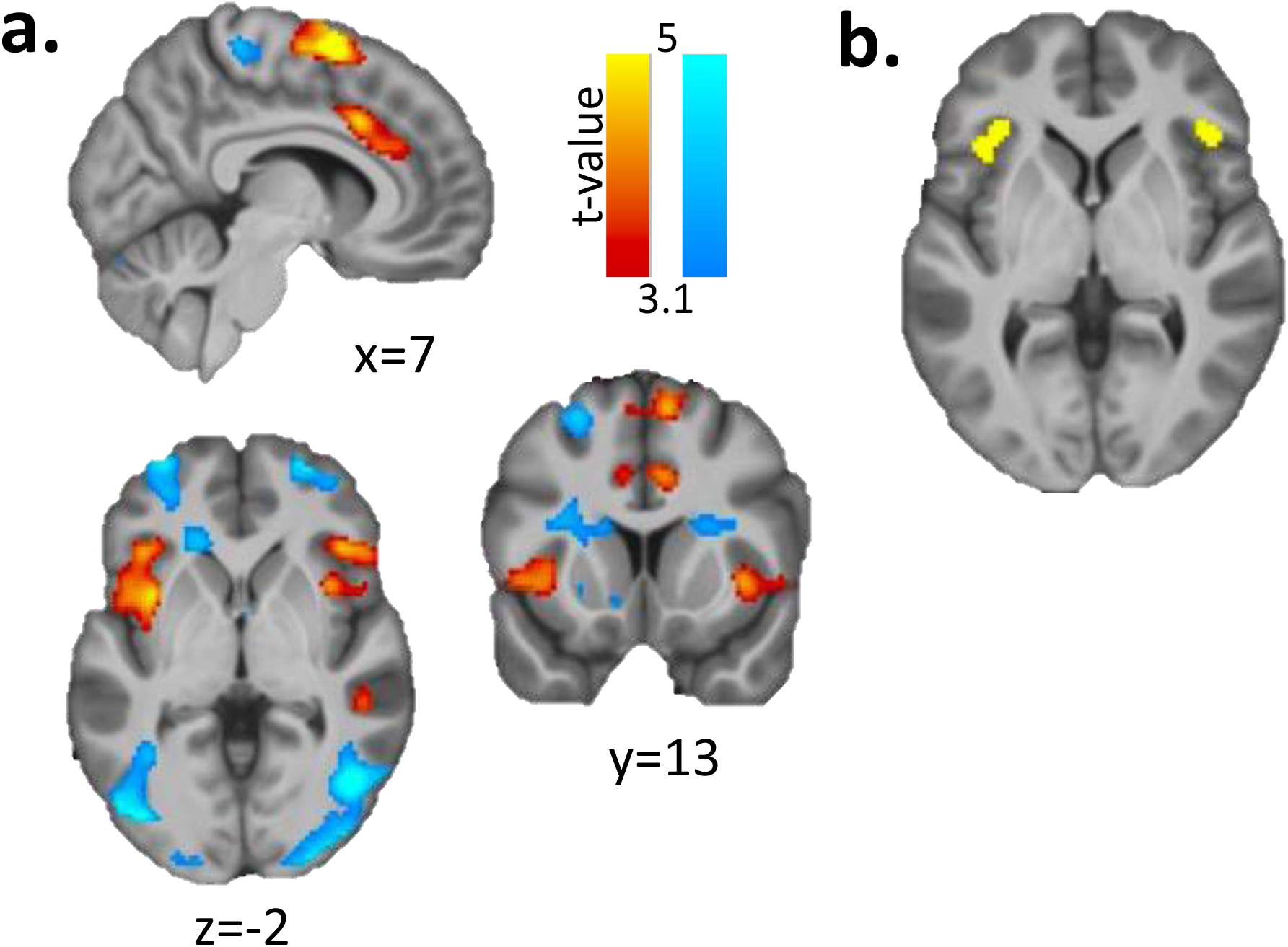
Neural correlates of slips of action after outcome-devaluation *Note*. Neural correlates of slips of action in the test phase, as revealed by increased (red – yellow) and decreased (dark – light blue) activity during devalued slips compared to upvalued responses. Clusters that survived whole-brain familywise error correction include increased activity in a fronto-parietal network, including the left anterior insula extending to the inferior lateral prefrontal cortex, supplementary motor area, dorsal anterior cingulate cortex, bilateral inferior parietal lobule and supramarginal gyrus. Conversely, lower activity was seen in the left anterior cingulate cortex extending into caudate nucleus, premotor/primary motor cortex, left lateral OFC, bilateral superior parietal lobe and several occipital/primary visual areas. Results are shown here at p<.001 (uncorrected) for visual purposes, overlaid on the mean T1 image of all participants. **B**. The bilateral anterior insula was found to be commonly activated during devalued slips (x = ±40, y = 26, z = 2). Shown here in yellow are the voxels that overlap between all four contrasts comparing devalued slips relative to correct (non-)responses during still-valuable, still-not-valuable, devalued and upvalued trials (thresholded at p<0.001 uncorrected).

As preregistered, we also compared upvalued NoGo responses (‘inhibition slips’) to correct devalued (NoGo) trials, but this did not reveal any significant activation patterns. Moreover, we were not able to conduct the contrast between upvalued and still-valuable NoGo trials, due to the low number of omission errors on still-valuable trials.

Our results thus identify the anterior insula as a common region associated with slips towards devalued outcomes, as activity in this region was higher during slips than during Go responses towards upvalued and still-valuable outcomes. However, both contrasts are confounded by expected value (the outcome value during the test phase) as they both compare stimuli signaling a non-valuable outcome (devalued) with stimuli signaling a valuable outcome (upvalued or still-valuable). To control for this, we ran some additional exploratory analyses, comparing activity during devalued slips with correct NoGo responses on devalued and still-not*-*valuable trials. Although these contrasts are difficult to interpret by themselves – they are themselves confounded by pressing a button or not – looking at the overlap between all four contrasts overcomes the value-related confounds and hence could find a common process in the expression of habits. To this end, we used ImCalc to create binary images of all four contrasts thresholded at t=3.1 (equivalent to p<0.001 uncorrected) and multiply them. The result of this exclusion analysis, which is akin but to a conjunction analysis, shows that the bilateral anterior insula was commonly activated across all four contrasts (Figure 4B).

##### Symmetrical outcome-revaluation test: comparing goal and implementation intentions

None of the planned contrasts comparing test-phase trials trained with implementation with goal intentions revealed significant activation patterns.

**Table 2.**
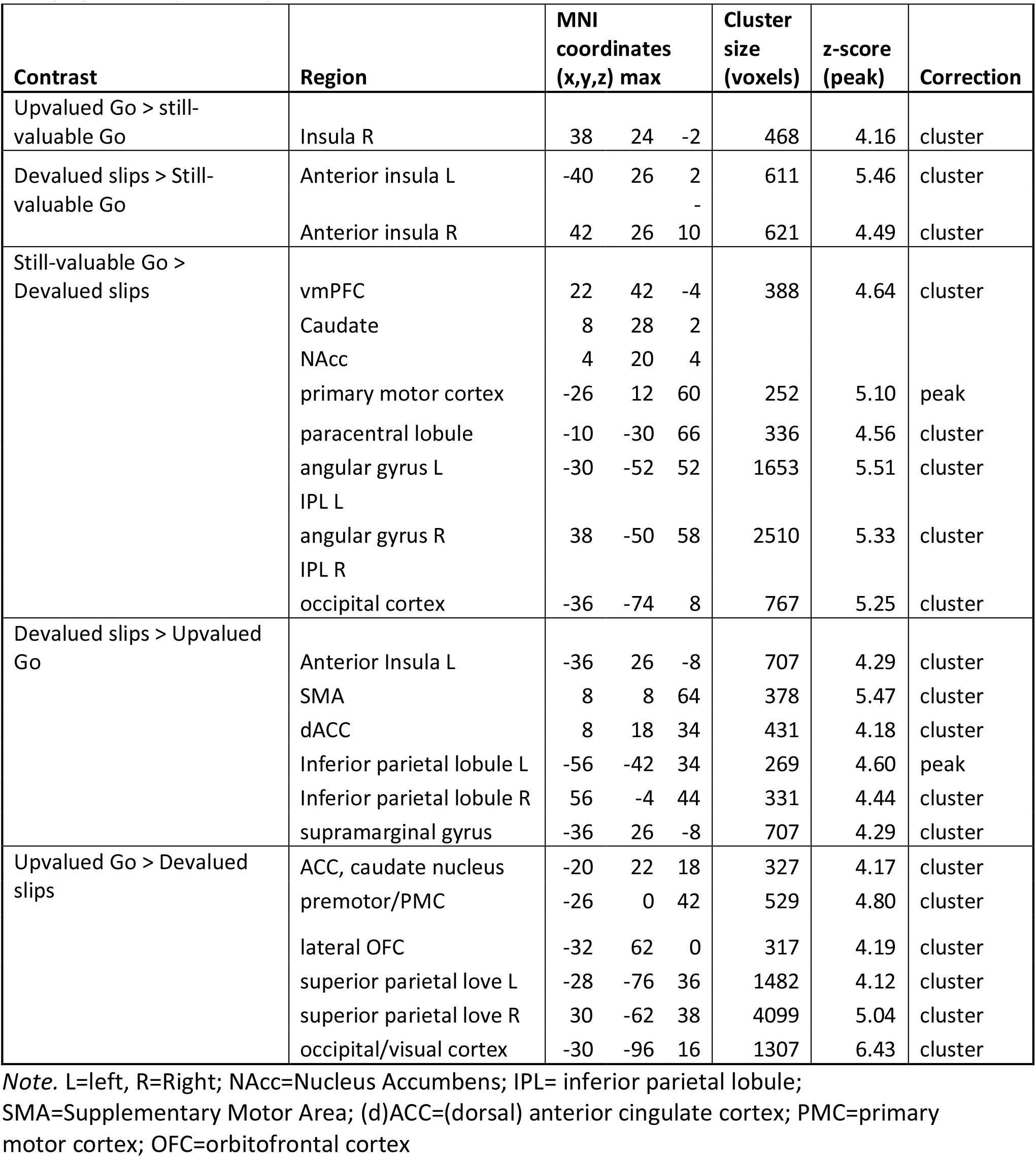
Imaging results of the test phase

## DISCUSSION

The aim of the present study was to investigate whether the brain can strategically go on automatic pilot. We investigated this by measuring the impact of strategic planning (i.e., implementation intentions versus goal intentions) on the acquisition of instrumental actions as well as subsequent flexible, behavioral adjustment. When strategic planning was first introduced during the instrumental learning phase of our paradigm, implementation intentions improved performance relative to goal intentions. Furthermore, in line with the idea that their beneficial effect was mediated by accelerated S-R learning, implementation intentions were associated with reduced activity in the anterior caudate, a brain area previously implicated in goal-directed control (Liljeholm et al., 2011; Watson et al., 2018).. These effects of strategic planning on performance and neural activity were only apparent early in training, with participants reaching high levels of accuracy (and reduced activity in the anterior caudate) by the end of the learning phase independent of intention type. Our central question, however, was whether implementation intentions would actually *impede* performance when flexible, behavioral adjustment was required during the subsequent outcome-revaluation test. Importantly, we found no evidence for a detrimental effect of strategic planning on the ability to adapt behavior to changing outcome values, nor any effect on underlying neural activity patterns. We conclude that strategic planning of S-R mappings may allow people to go on automatic pilot to increase behavioral efficiency, but that this does not have to come at the expense of behavioral flexibility. Therefore, mental rehearsal of S-R links does not appear to suffice for the formation of a rigid habit, refuting the notion of ‘instant habits’ and suggesting that behavioral repetition may be crucial for the development of rigid habits.

To shed light on the implications of these findings, we will first discuss them in some more detail, including the basic results (i.e., across strategies) on the relatively novel symmetrical outcome-revaluation task (SORT). Firstly, during the instrumental learning phase, we observed increasing accuracy and decreasing reaction times over the course of training, suggesting that participants acquired the S-R mappings. This interpretation was further supported by high levels of subjective automaticity of responding at the end of the instrumental learning phase and increasing involvement of two distinct parts of the dorsal striatum: the posterior putamen, replicating findings from Tricomi et al. (2009), and the caudate nucleus head. Several previous fMRI studies have indirectly implicated the dorsal striatum in habit learning, either showing that with longer instrumental training this region becomes more active (Tricomi et al., 2009; Wunderlich et al., 2012) or that functional connectivity with the (pre-)motor cortex increases (Horga et al., 2015; Zwosta et al., 2018). Although increased activity of the posterior putamen was only significant with small-volume correction (so not very robust), activity of the caudate nucleus as well as the hippocampus survived whole brain correction. Both regions have previously been implicated in the encoding of S-R representations (McNamee et al., 2015). Moreover, we found *de*creasing activity of the primary motor cortex (extending to mid-posterior cingulate), the hippocampus (extending into the putamen), bilateral temporal cortex, and the right anterior caudate, previously implicated in goal-directed control (Balleine & O’Doherty, 2010; Liljeholm et al., 2011). In line with previous findings, these results suggest that dissociable neural regions support instrumental learning. Notably, these findings were specific to learning to make Go responses for valuable outcomes. In contrast, we did not see any changes in neural activity over the course of NoGo training, despite high accuracy and reported automaticity at the end of training. Thus, our neuroimaging analyses do not provide evidence for the development of ‘inhibition habits’ (Jahanshahi et al., 2015).

Importantly, when strategic planning was introduced after the first twelve training blocks, implementation intentions initially improved Go performance (reflected by higher accuracy during the first block-set than in the preceding (pre-)training block-set), while goal intentions impaired it (reflected by lower accuracy and slower reaction times). In contrast, NoGo learning (i.e. withholding a response for non-valuable stimuli) was not affected by planning strategy. These findings replicate our previous results (van Timmeren & de Wit, 2022). Furthermore, in line with the notion of ‘instant habits’, this behavioral effect of implementation intentions early in training was accompanied by reduced activity in the anterior caudate relative to goal intentions. This early effect of implementation intentions quickly disappeared, however, and no differences with goal intentions were observed on accuracy and reaction times in later training blocks, nor on subjective automaticity after training. Therefore, in support of the notion of strategic automaticity, it appears that instrumental acquisition initially benefitted from if-then planning, while dependency on goal-directed control (as reflected by anterior caudate activity) was reduced.

So far, there have been very few neuroimaging studies that have compared the use of implementation and goal intentions to support behavioral performance. One study (Gilbert et al., 2009) showed that implementation intentions engaged the medial BA10 more (and lateral BA10 less) than a control condition, which was argued to reflect increased cue monitoring (and reduced internal information processing). This contrasts with our finding that implementation intentions lead to reduced engagement of the anterior caudate during the instrumental learning phase. However, their control condition was very different to ours. Whereas they specified the cue and the outcome that it signaled to be available (i.e., the S-O contingency; “if the cue appears, then I can score 5points) we used a goal intention control condition that specified the R-O contingency, which is arguably more akin to a typical goal intention (e.g., “I will exercise to lose weight”).

In the next phase of the SORT, signalled outcome values changed, requiring flexible adaptation of responding to the discriminative stimuli. This allowed us to determine whether strategic planning (during training) would induce the rigidity that is commonly regarded as a hallmark of learnt habits that are stamped in through behavioral repetition. However, we failed to find convincing evidence that if-then planning impaired the ability to flexibly adjust responding when signalled outcome values changed. This was despite the fact that participants struggled to adjust learned S-R mappings overall, as reflected in a strong main effect of congruency. Furthermore, in line with the behavioral findings, we also found no evidence for an impact of planning on neural activation patterns during the extinction test phase. Therefore, this first neuroimaging investigation of the effect of implementation intentions on behavioral flexibility in an outcome-revaluation paradigm failed to provide evidence for a shift from goal-directed towards habitual control.

The evidence for intact behavioral flexibility despite if-then planning contrasts with results from an earlier study with this paradigm (van Timmeren & de Wit, 2022). In that study, we found that implementation intentions impaired test-phase performance overall, but this did not lead to inflexibility as would be reflected by lower accuracy on incongruent trials specifically. This general impairment was most likely due to the fact that implementation intentions, by focusing attention on the S-R mappings, blocked learning about the S-O contingencies. To prevent this from happening in the present study, we altered our paradigm to promote active S-O learning at first training phase, before intentions were introduced. As a result, participants already acquired high levels of S-O knowledge when they started using strategic planning. Integrating findings from both studies, it appears that when the agent has full knowledge of the (S-O) contingencies implementation intentions do not impair flexibility. This finding is encouraging, because in most applied situations in real life, agents are perfectly aware of the three-term instrumental contingencies. Therefore, our results are in line with the idea of implementation intentions being ‘flexibly tenacious’ (Gollwitzer et al., 2008; Legrand et al., 2017): people benefit from if-then planning when the situation specified in their plan is encountered (here in terms of higher accuracy and lower reaction times during training), but are goal-directed in the sense that they only act on these planned S-R mappings when the signalled outcome is currently a goal.

Across intentions, however, we found that action slips towards devalued outcomes were associated with increased bilateral insula (both when compared to still-valuable and upvalued responses), replicating findings from the only study to date looking at neural activity during slips of action (Watson et al., 2018). The insula is a functionally heterogeneous region (Uddin et al., 2017) but the anterior part has been critically implicated in inhibition and error and salience processing (Chang et al., 2013; Uddin, 2015). Specifically, previous work shows that failure to inhibit a learned response (stop signal paradigm) is associated with bilateral insular activity (Ramautar et al., 2006). Additionally, when compared to upvalued responses, slips were associated with increased activity in the dorsal anterior cingulate cortex, the SMA and parietal cortex, all part of the salience network (Seeley et al., 2007). Conversely, lower activity during slips was seen in the vmPFC, or medial OFC, when compared to responses for still-valuable outcomes. A previous outcome-devaluation study suggested that activity in this region mediates goal-directed instrumental learning (Valentin et al., 2007). A similar contrast, comparing devalued action slips with responses towards upvalued outcomes, showed lower activity in the lateral OFC and the ACC/caudate nucleus head, regions that have also been implicated in goal-directed control. Overall, our results suggest that habitual slips of action arise as a consequence of lapses in goal-directed control (as reflected by decreased activity in these regions) rather than by increased activation of S-R habit regions (i.e., the dorsal striatum). Finally, the informal conjunction analysis of devalued slips (Figure 4B), controlling for differences in expected value and motor response, showed that the anterior insula was commonly activated across all contrasts, implicating it as a key region mediating habitual action slips.

A lack of reliable, positive markers of habits is an important issue in human habit research (De Houwer, 2019; De Houwer et al., 2018; Kruglanski & Szumowska, 2020; Watson, O’Callaghan, et al., 2022; Watson & de Wit, 2018). In the context of the present study, it begs the question whether habit strength independently contributes to stimulus-dependent, outcome-insensitive responding (i.e, slips of action). A recently published study with the SORT adds weight to this concern, as we showed there that extensive instrumental training failed to impair test performance (Watson, Gladwin, et al., 2022). The lack of reliable evidence for overtraining effects (see also de Wit et al., 2018) could mean different things, but our current findings may offer an interesting explanation. Specifically, we observed that when the planning manipulation was first introduced during training, not only did implementation intentions improve performance, but goal intentions also significantly *impaired* performance. This may indicate that participants’ spontaneous strategy up to that point had not been to form goal intentions, but instead to switch as soon as they could to the more efficient strategy of focusing on the S-R mappings. In other words, they may have spontaneously formed implementation intentions (Bieleke & Keller, 2021). Therefore, rather than improving their performance with the explicit implementation intention manipulation, we impaired it in the goal intention condition. Such an early shift to reliance on S-R associations (i.e., within twelve blocks of training) may explain that previous experimental studies failed to find evidence for overtraining, as their short training conditions may already have been sufficiently long to induce this, and beyond that early shift additional training may not have significantly enhanced the strength of those associations. This idea accords well with results from a study by Pool et al. (2022) who found that, following outcome devaluation on a free-operant task, already after moderate training (twelve blocks) outcome-insensitive habitual responding was seen in the majority of participants. Our findings further reinforce this interpretation by showing significant changes in neural activity over the course of this relatively short training, with activity of the anterior caudate (implicated in goal-directed learning) decreasing and of the dorsal striatum (implicated in habitual control) increasing. From our study it’s unclear, however, how activity in these regions developed in the earliest stages of instrumental training, as that took place outside the scanner. Future research should determine how many behavioral repetitions it takes to permit this shift to an S-R strategy, by assessing the effect of a goal intention manipulation at different time points during training. Our hypothesis is that at the start of training this would not yet have a negative impact – relative to implementation intentions – but that it will after a few blocks.

In conclusion, we provide evidence for increased efficiency but preserved flexibility following strategic if-then planning. These behavioral findings were mirrored in our analyses of the underlying brain activity: implementation intentions did not reduce the engagement of goal-directed control when goals changed, nor increase activity in habit regions. Therefore, our findings suggest that this strategic planning technique supports the implementation of a new target behavior while still allowing for flexible adjustment when goals change.

## Data availability statement

Data to recreate the main behavioral analyses and the full analysis pipeline with output (in JASP) are available at https://osf.io/yrpxa. Whole-brain t-maps (without thresholding) of the main fMRI contrasts are available at https://neurovault.org/collections/13191/.

## Author Contribution

Tim van Timmeren: Conceptualization; Data curation; Formal Analysis; Investigation; Methodology; Project administration; Visualization; Writing—Original draft; Writing—Review & editing. Nadza Dzinalija: Investigation; Project administration; Writing—Review & editing. John O’Doherty: Conceptualization; Writing—Review & editing. Sanne de Wit: Conceptualization; Funding Acquisition; Resources; Supervision; Writing—Review & editing.

